# Hippocampal subfields revealed through unfolding and unsupervised clustering of laminar and morphological features in 3D BigBrain

**DOI:** 10.1101/599571

**Authors:** J. DeKraker, J.C. Lau, K.M. Ferko, A.R. Khan, S. Köhler

## Abstract

The internal structure of the human hippocampus is challenging to map using histology or neuroimaging due to its complex archicortical folding. Here, we aimed to overcome this challenge using a unique combination of three methods. First, we leveraged a histological dataset with unprecedented 3D coverage, 3D BigBrain. Second, we imposed a computational unfolding framework that respects the topological continuity of hippocampal subfields, which are traditionally defined by laminar composition. Third, we adapted neocortical parcellation techniques to map the hippocampus with respect to not only laminar but also morphological features. Unsupervised clustering of these features revealed subdivisions that closely resemble ground-truth manual subfield segmentations. Critically, we also show that morphological features alone are sufficient to derive most hippocampal subfield boundaries. Moreover, some features showed differences within subfields along the hippocampal longitudinal axis. Our findings highlight new characteristics of internal hippocampal structure, and offer new avenues for its characterization with in-vivo neuroimaging.

## Introduction

The hippocampus is one of the most heavily investigated brain structures in neuroscience. Much research in recent years has focused on questions about its subdivisions, guided by the idea that different regions within the hippocampus may perform different functions and may also be differentially prone to disease (Small et al., 2011). These developments pose central question as to how to characterize subdivisions in anatomical terms. Traditionally, most proposed subdivisions have relied on histology and cytoarchitecture, leading to the notion of distinct hippocampal subfields that typically include the the subicular complex, Cornu Ammonis (CA) 1 to 4, and the dentate gyrus (DG) (see Duvernoy et al., 2013 for review). More recently, increasing interest has also emerged concerning graded differences along the anterior-posterior axis based on subfield composition and connectivity (Strange et al., 2014; Poppenk et al., 2013; Plachti et al., 2019). An organizational principle that shapes these dimensions, i.e., subfields and anterior-posterior differences, is the complex topology within the hippocampus that results from its ontological development (Duvernoy et al., 2013; DeKraker et al., 2018). This principle has received only limited investigation to date but requires careful consideration in any effort to characterize the internal architecture of the hippocampus. The current paper aims to investigate the relationship between hippocampal topology, morphology, and cytoarchitecture in humans, taking advantage of the unique and powerful “Big Brain” dataset that provides continuous histological sampling with full 3D coverage (Amunts et al., 2013). A particular promise of this approach lies in its applicability to in-vivo Magnetic Resonance Imaging (MRI).

While commonly used MRI measures do not allow for cytoarchitectural characterization, MR-based protocols have been developed to indirectly infer the locations of hippocampal subfields in humans based either on manually delineated landmarks or corresponding probabilistic atlases that are informed by histological reference material (Yushkevich et al., 2015; Yushkevich et al., 2015; Iglesias et al., 2015). However, traditional histological references can be problematic for several reasons. First, they often contain only select coronal slices taken from regions where folding is simplest, most frequently from the hippocampal body (with the notable exception of Ding & Van Hoesen, 2015). Second, even in the hippocampal body slices are taken sparsely, limiting the amount of contextual features that can be gathered from neighboring slices or other planes of view. Third, histological preparation often deforms the tissue of interest relative to its in-vivo state, which is a problem for MRI co-registration unless the histological sample is also imaged prior to histological preparation. Finally, even among neuroanatomists there is some disagreement as to exactly which labels, stains, and histological features should be used for defining hippocampal subfields (see Wisse et al., 2017). Some previous studies have made use of ex-vivo MRI to aid in the translation of histology to MRI (Yushkevich et al., 2009; Iglesias et al., 2015) in an effort to mitigate some of these issues. However, even with such an approach, inter-individual differences in hippocampal morphology can impose limitations for inferring subfields or other structural features, when hippocampal topology is not considered.

It is well established that the human hippocampus is a folded component of archicortex that is continuous with the neocortex (as summarized, for example, by Duvernoy et al., 2013; Nieuwenhuys et al., 2013). The hippocampal folds include wrapping around its innermost region – the DG, as well as anterior-posterior folding that is sometimes referred to as dentation, digitation, or gyrification. The gyrification seen in the hippocampus is morphologically similar to gyrification in the neocortex (although not necessarily based on the same ontogeny). It has been shown to vary considerably between individuals (DeKraker et al., 2018; Chang et al., 2018) and can be correlated with age (Cai et al., 2019) or disease, such as temporal-lobe epilepsy (Blümcke et al., 2013). This folding is an important aspect of understanding the internal structure of the hippocampus, and for appreciation of the continuity of subfields, particularly in its anterior portion that includes the uncus (Ding & Van Hoesen, 2015). Topological analyses can provide a framework for extracting these continuities, for example through unfolding (DeKraker et al., 2018), and offer the basis for laminar and further morphological characterization of complete hippocampal structure in 3D, including subject-specific gyrification.

The dataset made publicly available by BigBrain (Amunts et al., 2013) provides a unique opportunity to conduct topological analyses of histology data in 3D, and to examine topological measures in unfolded tissue. This dataset consists of 3D histology, digitally reconstructed from images of a serially sectioned and stained healthy cadaveric brain. In the current project, we used reconstructed blocks of the left and right hippocampi (40um isotropic) to identify topologically-derived laminar and morphological features under our hippocampal unfolding framework. To characterize laminae, we focused on 10 computationally derived features describing the distributions of neurons (Amunts et al., 1999). Morphological features were also computationally derived and included thickness, curvature, inner and outer surface textures, as well as gyrification. We then compared these morphological and laminar features to classic descriptions of subfields and examined variations along the anterior-posterior hippocampal axis. We anticipated that the features examined would differ substantially between subfields. Therefore, we also tested whether it might even be possible to obtain successful subfield segmentation with an unsupervised feature-based approach. This type of approach is desirable for is objectivity, which could help resolve differences among neuroimagers and histologists on subfields definitions. It also allows us to examine which subsets of features are sufficient to derive clusters resembling ground truth hippocampal subfields. For this purpose, we contrast the contributions of laminar and morphological features, given that laminar features are used most prominently in histology (see Duvernoy et al., 2013; Nieuwenhuys et al., 2013) but morphological features, such as thickness, are more readily available in high-resolution structural MRI (e.g. DeKraker et al., 2018).

## Results

The backbone of our analyses was to impose a topological unfolding framework to manual hippocampal traces, a method that we previously developed for 7T MRI (DeKraker et al., 2018). We then extracted various morphological features of hippocampal structure from the left and right 3D BigBrain hippocampi. We computed laminar features based on the work of (Amunts et al., 1999) and modeled as in (Waehnert et al., 2014). We then performed unsupervised, data-driven clustering of these features and compared resulting clusters to manually segmented hippocampal subfields. Finally, we examined differences in hippocampal structure along its longitudinal (i.e., anterior-posterior) axis.

### Manual Tracing

Figure 1 shows BigBrain coronal slices alongside manually segmented subfields in the head, body, and tail of the hippocampus, as well as corresponding 3D models. Several features were detected in tracings of the hippocampus in 3D BigBrain that were not detected in previous in-vivo MRI work that we know of. Clusters of pyramidal cells or ‘islands’ can be seen on the inner surface of the subiculum (stratum lacunosum), which have been observed in histology throughout the presubiculum (Duvernoy et al., 2013; Ding & Van Hoesen, 2015) and others). A medial and anterior fold along the vertical component of the uncus, approximately 0.3mm thick and up to 3.6mm in length, was observed, as described in (Duvernoy et al., 2013; Ding & Van Hoesen, 2015). Finally, numerous gyrifications throughout the posterior body and tail of the hippocampus were observed, which was also observed using MRI in (Chang et al., 2018), though not to the extent seen here. This was most prominent in CA1, but was also present in the DG and in CA4 which followed the same gyrification scheme as CA1. Models of the DG alone and additional anatomical notes can be found in Supplementary Materials section A. Total volumes of each subfield can be seen in Table 1. Note that these volumes are smaller than what is typically reported in MRI, which may be due to our exclusion of alveus and SRLM laminae which can be hard to differentiate from partial voluming in MRI, but may also be influenced by tissue shrinkage during histological processing.

**Table 1.**
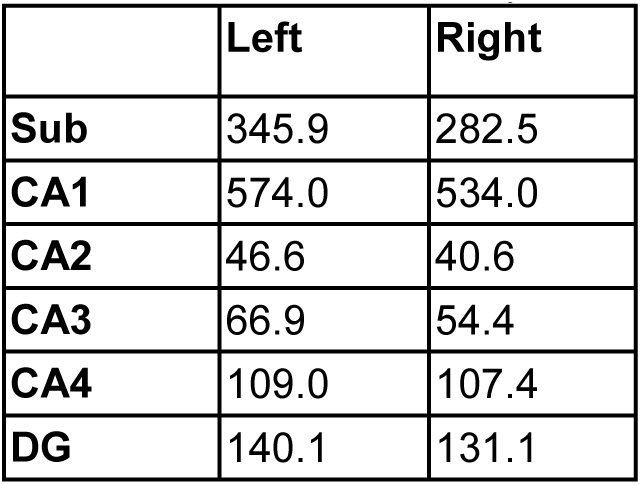
Volumes of each manually defined subfield (mm^3^).

|  | Left | Right |
| --- | --- | --- |
| **Sub** | 345.9 | 282.5 |
| **CA1** | 574.0 | 534.0 |
| **CA2** | 46.6 | 40.6 |
| **CA3** | 66.9 | 54.4 |
| **CA4** | 109.0 | 107.4 |
| **DG** | 140.1 | 131.1 |

**Figure 1.**
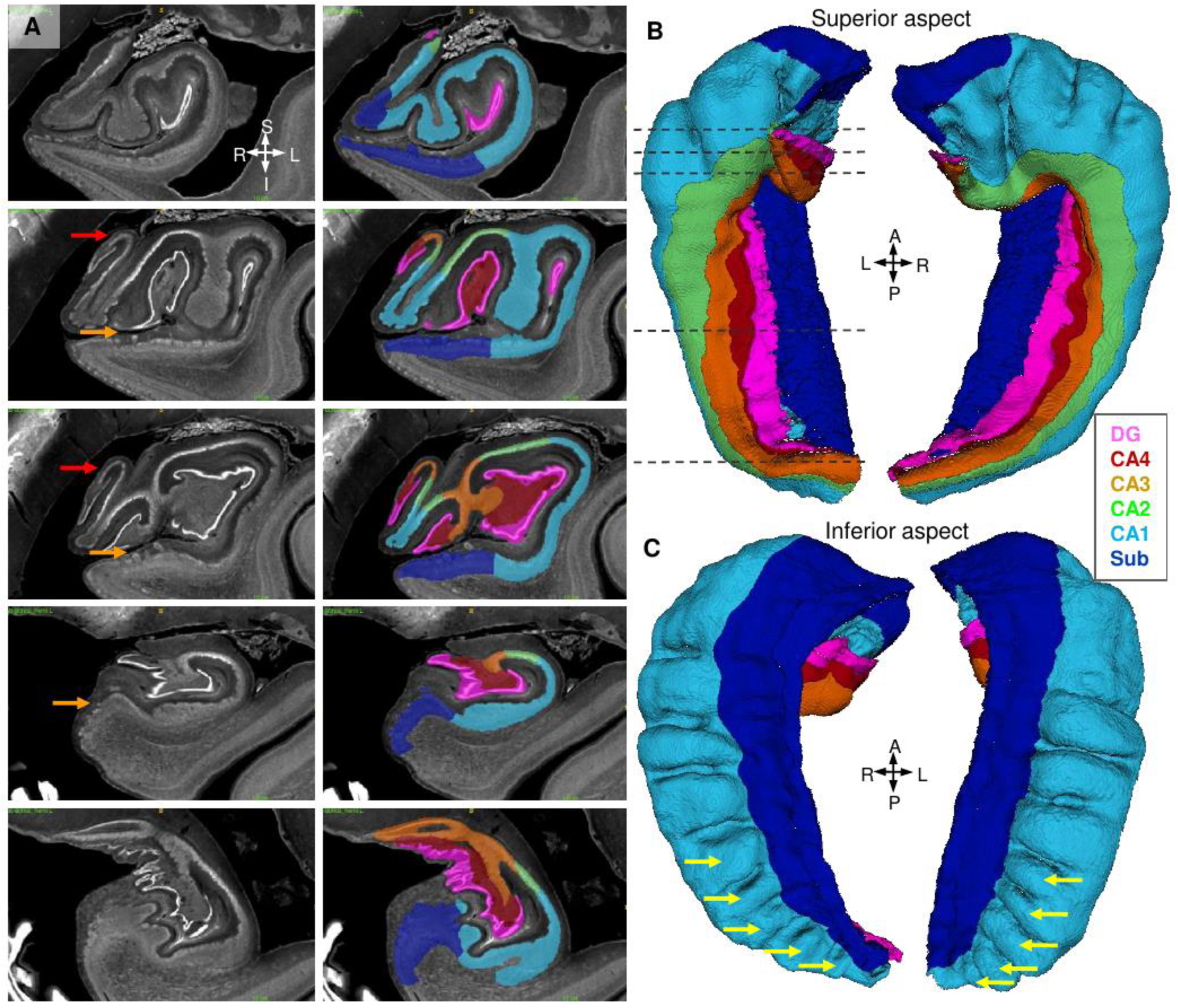
Manual traces of hippocampal archicortex and segmentation into subfields. A) shows coronal slices through the left hippocampal head (rows 1-3) body (row 4) and tail (row 5), with manual segmentations overlaid in the images to the right. B) shows 3D models of each hippocampus as seen from their superior aspect, with the inferior aspect shown in C). Dotted lines in B) indicate approximate locations of each coronal slice shown in A). SRLM, vestigial hippocampal sulcus, alveus, and fimbria were excluded from all labels. Red arrows indicate anterior folding in the vertical component of the uncus, orange arrows indicate ‘islands’ of neuronal cell bodies in the subicular stratum lacunosum, and yellow arrows indicate gyrifications in the posterior body and tail of the hippocampus.

### Topological unfolding

Figure 2A shows the proximal-distal and anterior-posterior Laplacian solutions which make up the two axes of our topological unfolded space. The DG was not unfolded. Although it was easily distinguishable from other subfields by its very high cell density it is topologically disconnected from the rest of the archicortex, and therefore would be out-of-plane (i.e. perpendicular) to our unfolded space (see Figure 1 for visualization). Figure 2B shows a mid-surface mesh of the hippocampus, coloured according to manually segmented subfields as in Figure 1. This surface was then mapped to 2D unfolded space according to the anterior-posterior and proximal-distal Laplace solutions. In unfolded space, subfields are relatively constant from anterior to posterior, with subiculum being proportionally larger in the very anterior and smaller in the very posterior, however these may be artefacts of manual segmentation since these region are very small in native space. This unfolding is illustrated in our online video (created through linear interpolation of all points between native and unfolded space).

**Figure 2.**
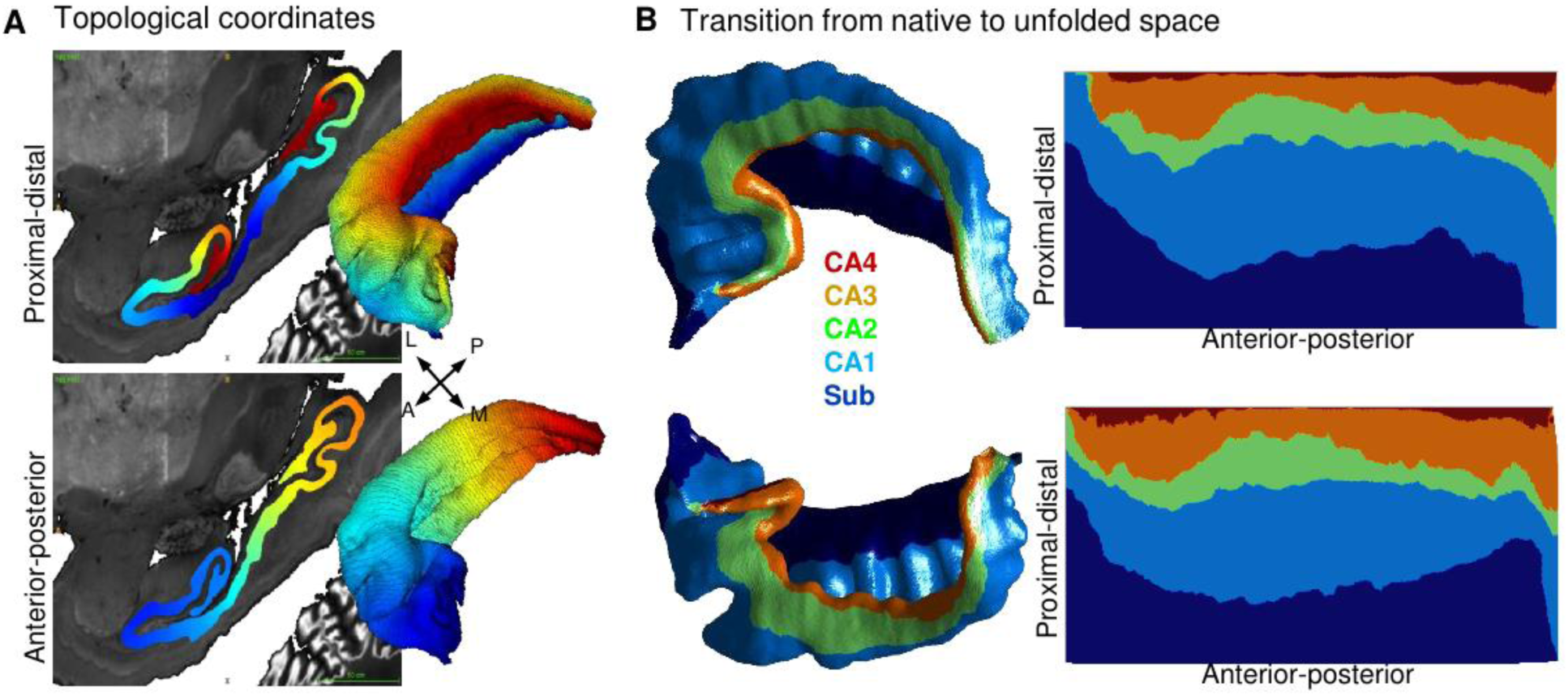
Topological unfolding framework in 3D BigBrain with hippocampal subfields. A) Sagittal slice and 3D models of the Laplacian solutions (proximal-distal and anterior-posterior) for the right hippocampus. B) Mid-surface topological models of the left and right hippocampi in native and unfolded space.

### Characterization of the hippocampus in unfolded space

Figure 3 shows a full characterization of the left and right BigBrain hippocampi with respect to the 5 morphological and 10 laminar features. These features are illustrated at the top of the figure, but additional details can be found in the Materials and Methods section. As in related work (Duvernoy et al., 2013; DeKraker et al., 2018), thickness was highest in the subiculum and CA4 and lowest in CA2. Curvature was generally high in subiculum, which reflects its outward curling away from the rest of the hippocampus, but in CA1 vertical bands of positive and negative values can be seen which correspond to the hippocampal gyrifications seen in Figure 1. This region is also highlighted by our gyrification measure, which differs from curvature in that it does not vary by direction. Inner surface texture shows an almost honeycomb texture that is most prominent in the subiculum where subicular ‘islands’ of neurons are found in stratum lacunosum (Duvernoy et al., 2013). Outer surface texture appears smoother, and more closely resembles the mid-surface curvature measure. Note that the surface textures measures differ from the curvature measure only in that they capture very local details. Thus, they may not be available in lower resolution data. Thus features such as thickness and gyrification may be especially of interest in translation of this work to MRI, particularly because they show such clear distinction between subfields.

**Figure 3.**
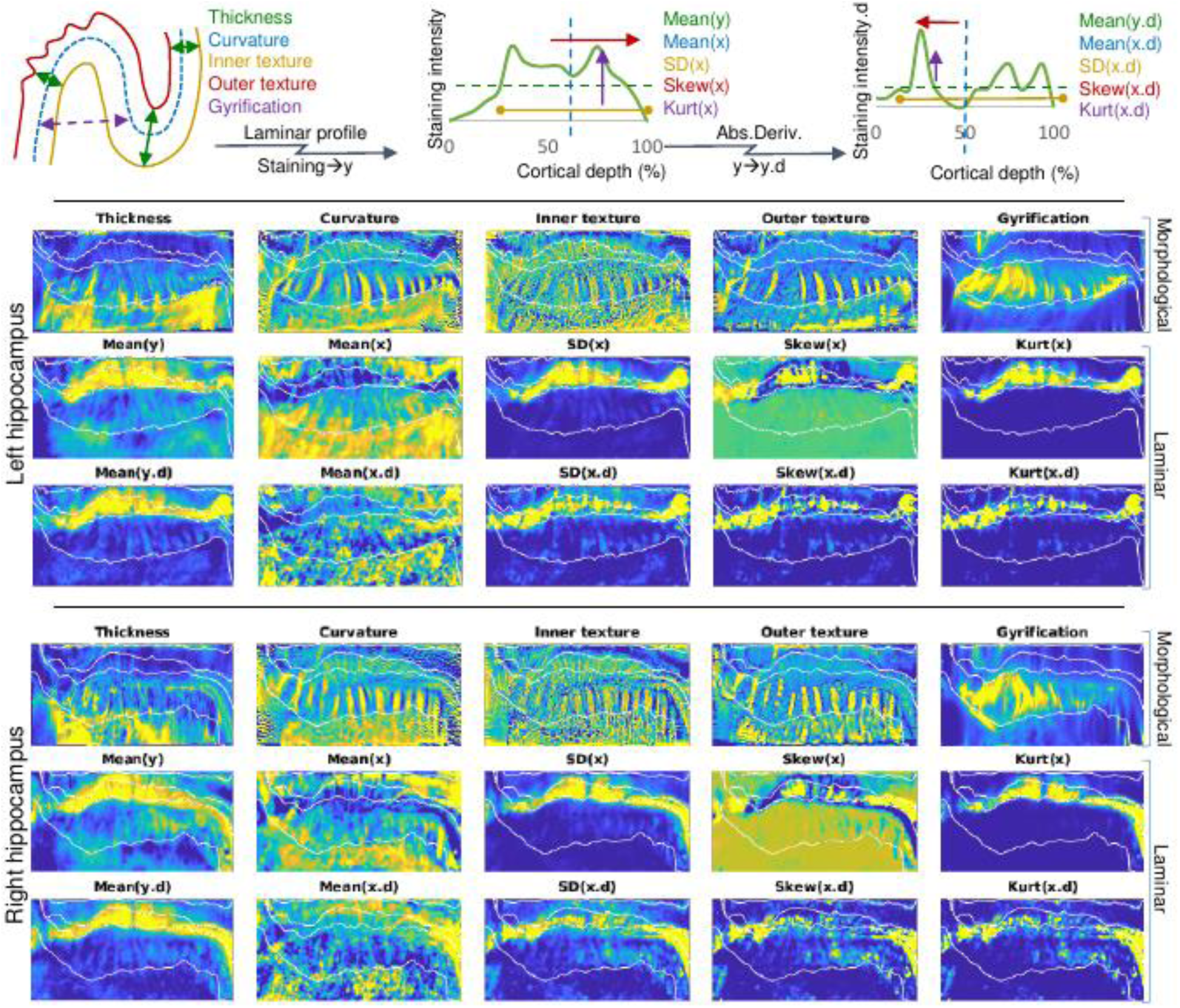
Characterization of the hippocampus using morphological and laminar features. The top diagrams illustrate how each feature is derived (see Materials and Methods section for details). Top left shows an example segment of cortex, while the top centre and top right show an example laminar profile and its absolute derivative (Abs.Deriv), respectively (Amunts et al., 1999). Heat maps below show the z-scored values of each feature across the unfolded hippocampus in the left and right hemispheres. Overlaid in white are manually defined subfield borders, the top edge being the border with the DG which is out-of-plane.

Of the laminar features computed here, Mean(y) was highest in region CA2, which also agrees with the high neuronal densities observed in this region (Duvernoy et al., 2013). Mean(x) showed almost the inverse pattern, with high values in all regions except CA2. This means that the distribution of neurons was shifted towards the inner surface in CA2. SD(x) was highest in CA2, indicating a wide distribution of neurons relative to the thickness of that tissue. This was counter-intuitive since in native space CA2 appears to have a tight distribution of neurons, however, relative to its small thickness the distribution is wide. The fact that the SD(x) and thickness features differ so drastically, despite our intuition from native space images that they should be similarly small in CA2. The remaining 7 laminar features become more complex and quite similar to Mean(y), Mean(x) or SD(x). Thus some of these features may be redundant but we nevertheless included them for consistency with previous work in the neocortex (Amunts et al., 1999).

### Unsupervised identification of hippocampal subfields using combination of morphological and laminar features

By visual inspection, many of the features in Figure 3 show a clear distinction between the different manually defined subfields. Therefore, we sought to determine whether a combination of these features could be used to derive some or all of the subfield boundaries between subiculum and CA1 to CA4 computationally, using PCA followed by k-means clustering (see Materials and Methods section for details). In this endeavor we also examined whether morphological or laminar features in isolation would be sufficient to allow for successful clustering, i.e. to derive clusters that closely resemble ground truth hippocampal subfields. For consideration of morphological features, we excluded surface textures given that they include subicular ‘islands’, which arguably also qualify as laminar features (see Materials and Methods section for further discussion). Figure 4 shows the results of unsupervised clustering of the combined feature sets, laminar features only, and morphological features only. We compared clusters to their closest corresponding manually defined subfield (ground truth) using Dice overlap scores in Table 2. When all features were combined in this analysis, good (0.7) to very good (0.8+) overlap was found for most subfields. Specifically, subfields subiculum, CA1, as well as combined CA2 and CA3 showed overlap with ground truth segmentations. Manually defined region CA2 had two clusters that overlapped with it (orange and green in Figure 4). The green cluster corresponded to the most dense regions of CA2 (e.g. where Mean(y) and SD(x) were high), and several other laminar features echoed this pattern. The fact that multiple features echoed this pattern may have contributed to why two clusters were generated in CA2 rather than just one. In other words, the variance within CA2 may have been amplified by the presence of redundant features. Using a combination of labels CA2 and CA3, as is often done in MRI segmentation protocols (Yushkevich et al., 2015), increased the Dice overlap scores as expected. We note that subfield CA4 did not emerge as a unique cluster and was instead included in the same cluster as CA1 or CA3. This remained true even when the number of clusters (k) was increased up to k=16 (Supplementary Materials section D). Overlap of CA4 with CA3 is to be expected given their topological closeness, but overlap with CA1 is more surprising. One possible explanation is that despite their topological separation, both of these regions were thicker and contained a lower density of neurons than the other CA fields. Finally, the current analyses did not reveal any evidence for the subregions of the subicular complex as described by (Ding 2013). This is not surprising because BigBrain only contains a single contrast (neuronal cell bodies); other contrasts (particularly myelin) or even immunochemical profiles are typically used to detect these subregions (Ding & Van Hoesen, 2015; Ding, 2013).

**Table 2.** Dice overlap scores between k-means clusters and their closest corresponding manually defined subfield.

|  | All features |  | Laminar features |  | Morphological features |  |
| --- | --- | --- | --- | --- | --- | --- |
|  | Left | Right | Left | Right | Left | Right |
| **Sub** | 0.92 | 0.91 | 0.81 | 0.80 | 0.92 | 0.86 |
| **CA1** | 0.74 | 0.73 | 0.59 | 0.66 | 0.80 | 0.74 |
| **CA2** | 0.73 | 0.66 | 0.72 | 0.64 | 0 | 0 |
| **CA3** | 0.56 | 0.58 | 0.49 | 0.45 | 0.59 | 0.56 |
| **CA2&CA3** | 0.83 | 0.78 | 0.78 | 0.70 | 0.84 | 0.80 |
| **CA4** | 0 | 0 | 0 | 0 | 0 | 0 |

**Figure 4.**
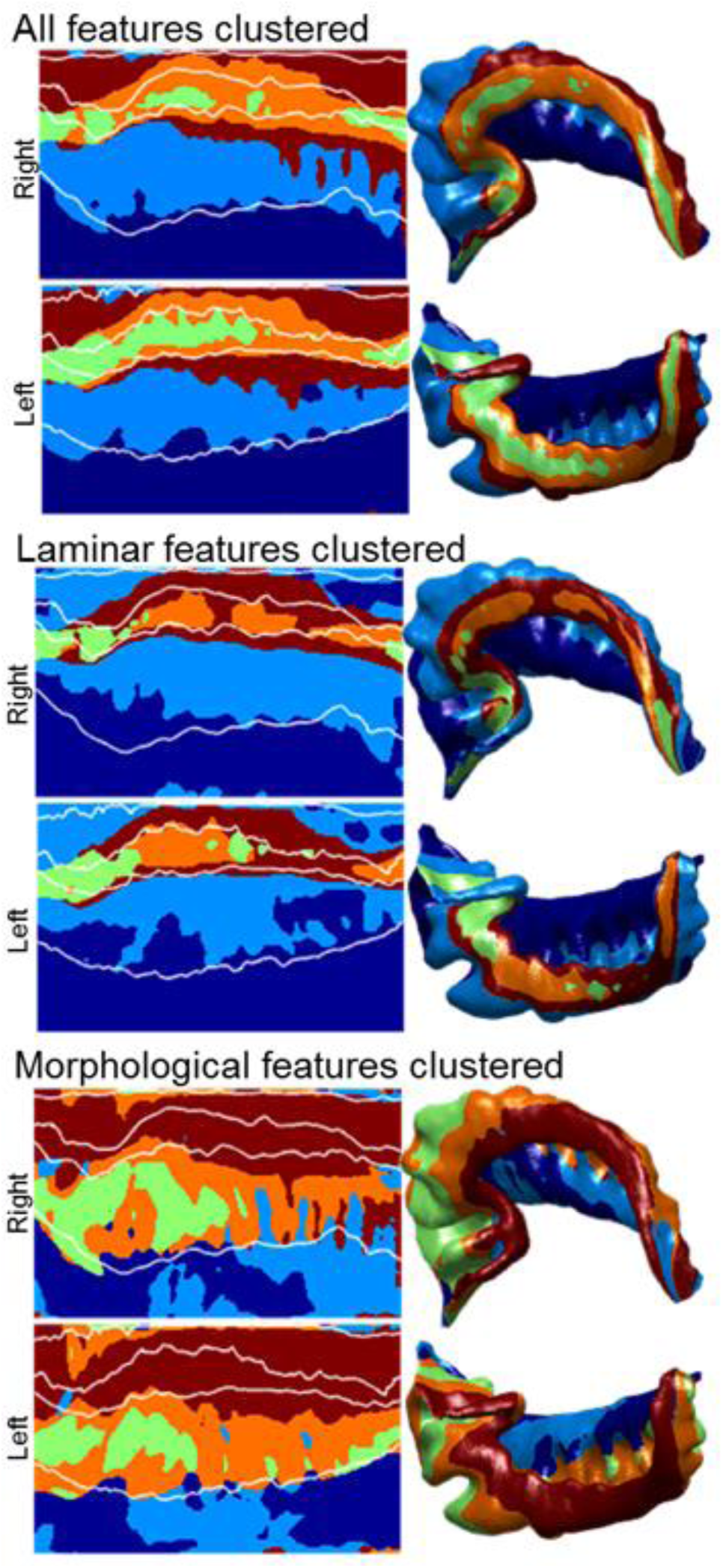
Unsupervised k-means clustering of features. The left images show k-means clusters in unfolded space with k=5, with manually defined subfield borders overlaid in white. The right images show the same data in native space with 10% anterior and posterior edges extrapolated by nearest neighbour. Clustering was completed for the combined set of all features, laminar features only, and morphological features only.

### Unsupervised identification of hippocampal subfields using morphological or laminar features in isolation

We next asked whether subsets of features (i.e. morphological features alone or laminar features alone) could be used to derive hippocampal subfield borders. Laminar features alone were able to capture most boundaries with good accuracy, with the exceptions of CA1, CA2, and CA3 which had Dice scores below 0.7. (Figure 4; Table 2). Again, combining CA2 and CA3 lead to good (0.7+) agreement with manually defined ground truth segmentations. CA1 was less well defined using only laminar features, and indeed there is some disagreement over the exact border between subiculum and CA1 in the histological literature (some disagreement may depend on the inclusion of prosubiculum as its own region or simply a transition zone; see Wisse et al., 2017). Morphological features alone revealed two clusters within subiculum and two within CA1, and did not differentiate between CA2 and CA3 at all. Clustering using these features also highlighted boundaries surrounding CA4, but CA4 did not contain a unique cluster. Rather, the same clusters that were assigned to CA1 were assigned to CA4, similarly to when all features were used in clustering. However, it is worth noting that when their topological separation is considered visually, CA4 can easily be distinguished from CA1. Overall with the exception of differentiating CA2 from CA3, morphological features were sufficient to delineate hippocampal subfields with very good (0.8+ in most cases) accuracies, at a level similar to clustering based on the combination of all features.

### Relative contributions of individual features to subfield clustering

In order to better understand the inherent structure of the data used in the above k-means clustering of all features, we revisited the PCA that guided clustering and examined various PCA metrics. Figure 5A shows the total variance explained by each PCA component, and only the first 7 components explained >1% of the variance and were included in subsequent analyses. Figure 5B and D show a breakdown of the first 7 principal components. The first and most prominent component was most highly correlated with most laminar features, except Mean(x), Skew(x), and Mean(x.d) which showed an anti-correlation. Visualization of this component shows consistently high values in CA2. This makes sense that most laminar features showed uniquely high values in CA2, while Mean(x), Skew(x), and Mean(x.d) contained low values in this region (Figure 3). Subsequent components explain a decreasing portion of the total variance in the data, but correlate with different input features. Visual inspection of these components shows that some loosely follow the contours of the subfields. For example, component 3 quite clearly alternates low and high between subiculum, CA1, CA2, and CA3. Others, particularly components 5-7, appear to contain little subfield-related variance and may reflect noise captured by the later components. Interestingly, components 2 and 3 appear to show gradual anterior-posterior differences, with lower values in the anterior and higher in the posterior in component 2 and vice-versa in component 3.

**Figure 5.**
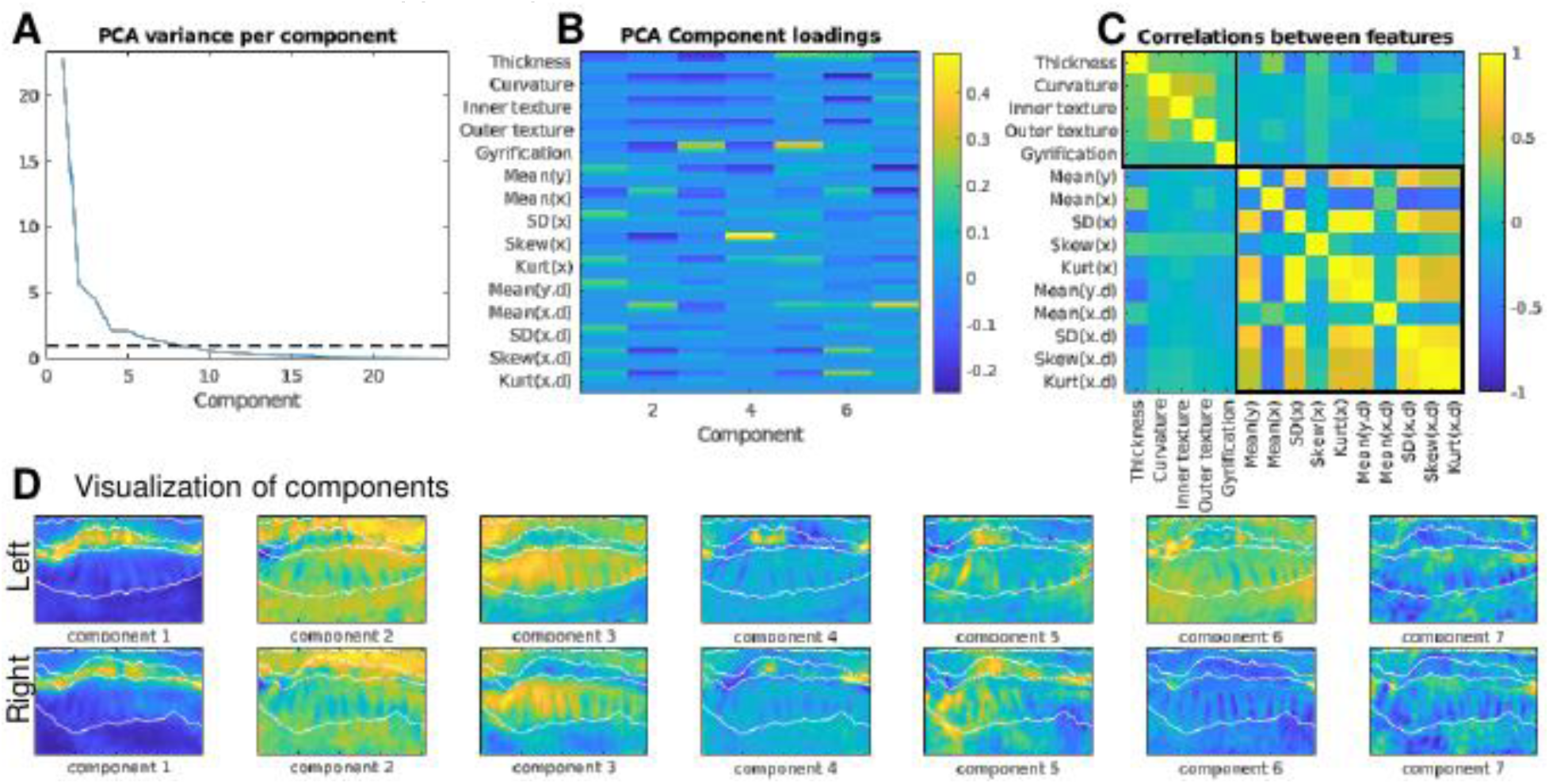
Exploration of inherent feature variance. A) shows PCA component loadings from each feature with a dotted line at 1% after which subsequent components were discarded. B) shows the loadings of the first 7 components on each feature, with multiple rows for the various smoothing kernels applied to each feature. C) shows the correlation between all features, with separate boxes around morphological and laminar features. D) shows a visualization of the first 7 components, with manually defined subfield borders overlaid in white.

Of the features used in this analysis, some were more correlated with each other than others (Figure 5C). In particular, all morphological features tended to be correlated with each other while all laminar features tended to be correlated or anti-correlated with each other, with only small correlations between morphological and laminar features. The fact that laminar features tended to be uncorrelated with morphological features is in line with the goal of the Equivolume model (Waehnert et al., 2014) which we applied in order to remove the effects of curvature on laminar displacement. Thus, overall, when modelled in 3D using the appropriate methods, morphological and laminar features represent different levels of structural information about tissue within the hippocampus.

### Structural variation along the longitudinal hippocampal axis

In a final set of analyses, we aimed to explore qualitatively whether subfields would show differences in feature composition along the anterior-posterior axis of the hippocampus. Towards this end we visualized possible trends along the axis in each manually-defined subfield (Figure 6). We primarily focused on the features gyrification, thickness, and mean neuronal density (Mean(y)) given that these features showed high contrast between different subfields. (Data for all other features are included in Supplementary Materials section A). Note that with this visualization, a high degree of separation can be seen between some subfields, as previously described (see Figure 3). Thickness and gyrification tended to show lower values at the anterior and posterior extremes, or in the vertical component of the uncus and tail of the hippocampus, which was also observed during manual tracing (Figure 1). However in the remainder of the hippocampus, namely the head and body, thickness remained relatively constant in each subfield while gyrification gradually decreased, as observed during manual tracing. Neuronal density was notably lower in most subfields in the anterior, approximately corresponding to the vertical component of the uncus, but showed only minor variations throughout the rest of the hippocampus. Overall, these visualizations suggest that anterior-posterior differences are most notable in gyrification.

**Figure 6.**
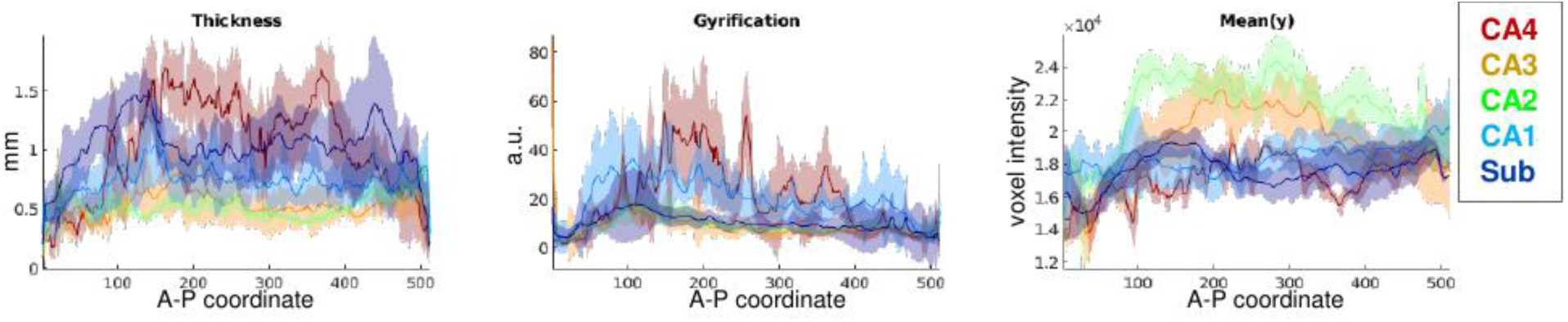
Features of interest plotted with respect to the anterior-posterior axis of the hippocampus. Colours indicate manually defined subfields, and shaded areas indicate standard deviation. Data are combined across the left and right hippocampi. a.u. stands for arbitrary units, see Materials and Methods section for additional details.

## Discussion

In the present study we show, for the first time, unsupervised clustering of human hippocampal subfields that closely resembles manually defined ground truth. We additionally show that morphological features alone are sufficient to derive most hippocampal subfield boundaries. Moreover, our findings reveal that some features, most notably gyrification in CA1, showed within-subfield differences along the anterior-posterior hippocampal axis. The current study sheds new light on the relationship between hippocampal topology, morphology, and laminar cytoarchitecture with respect to hippocampal subfields and the anterior-posterior axis.

### Structural characterization of the hippocampus in BigBrain

Manual tracing and 3D modelling of the hippocampus (Figure 1) at the level of resolution available in BigBrain revealed several features not seen in any 3D atlas that we are aware of. First, medial folding in the posterior end of the vertical component of the uncus was observed, similar to the inward ‘curling’ of the CA fields around the innermost DG in the rest of the hippocampus. Second, ‘islands’ of pyramidal neurons were present in stratum lacunosum in the subiculum. Third, gyrifications were present throughout the head, body, and tail of the hippocampus but were most prominent in CA1. These gyrifications were also echoed in the underlying DG (where the term dentation is often used to refer to this feature) and region CA4 that the DG partially encircles. Each of these features has been described in histology (Ding & Van Hoesen, 2015; Duvernoy et al., 2013), but has not been reconstructed in a 3D model at this level of detail. For example, (Adler et al., 2018) and (Iglesias et al., 2015) both performed detailed and fully 3D segmentation of the hippocampus and its subfields using ex-vivo MRI data, with additional histological data in the same participants provided by Adler et al., Our approach extends beyond these studies by utilizing higher-resolution tracing and by using histological cues inherent in the same images. Furthermore, our manual traces and quantitative analyses fully respect the topology of the hippocampus and, in turn, the continuity of each subfield throughout the entire length of the hippocampus. We note that the topology developed here does not cover the DG, which has its own topological arrangement that is perpendicular the rest of the cortex (including archi- and neo-cortex). This difference in topology arises from a different trajectory in ontogeny, in which the DG ‘breaks with’ the rest of the cortex and wraps around the distal-most archicortex, i.e., CA4 (Duvernoy et al., 2013; Nieuwenhuys et al., 2013). In future work, the DG could also be unfolded using a general framework similar to what is presented here, but critically, this approach would require employment of endpoints in a different plane.

After applying our topological unfolding framework, we computationally extracted morphological and laminar features from the hippocampus (Figure 3). Many of these features agree with qualitative descriptions by neuroanatomists, as discussed in the Results section. Some of these features may be informative for in-vivo imaging as well. For example, measures of thickness and gyrification can be obtained under our topological unfolding framework given sufficiently detailed segmentations, regardless of the availability of cytoarchitectonic features. These two features in particular show good contrast between subfields Subiculum, CA1, and CA4, and so they could be explicitly leveraged to guide segmentation or registration to histological reference materials in future MRI work. This may have been underappreciated in other in-vivo studies, including our own previous MRI study, where some of the gyrifications in the body and tail of the hippocampus could not be detected. This lack of detail would also lead to overinflated thickness measures, larger overall volumes, and perhaps differences in the proportional sizes of some subfields along the anterior-posterior extent of the hippocampus. Quantitative MR, such as T1 mapping, may additionally provide cues to approximate cytoarchitectonic features and indeed in our previous work we observed higher T1-weights in CA2 and CA3 (DeKraker et al., 2018) which may be driven by higher neuronal cell densities as observed in the current study. Thus the features described here show clear promise for characterizing or segmenting the hippocampus in future MRI work.

### Unsupervised clustering of all features reveals hippocampal subfields

We performed unsupervised clustering of all features to determine whether we could identify the classically described hippocampal subfields using a completely unsupervised computational approach. Results from clustering yielded generally high overlap with manual subfield segmentations in most cases (Table 2), with several exceptions that are outlined in the Results section. One particularly interesting observation was that CA4 was consistently assigned the same cluster as CA3 or CA1, even though it shares no topological boundary with CA1. The shared structural elements between CA4 and CA1, particularly their relatively high thickness, gyrification, and low density of neurons, may relate to why certain diseases, such as subtypes of epilepsy, selectively affect CA1 and CA4 similarly (Duvernoy et al., 2013; Blümcke et al., 2013). In future imaging work, CA1 and CA4 can be differentiated from each other, particularly under our unfolding framework, due to their topological separation.

Finally, to further explore the inherent dimensionality of these structural features, we examined the principle components of all features (Figure 5). From this we noted that the most prominent components varied in such a way that followed the contours of some or all subfield borders (see Results section). This suggest that the inherent structural variance in the hippocampus most naturally follows a proximal-distal patterning as seen in the classic subfield definitions. Some components additionally hinted at inherent anterior-posterior differences across the hippocampus.

### Morphological features are sufficient to approximate most subfield boundaries

In addition to clustering using all features, we also asked whether hippocampal subfields could be derived using only the subset of morphological features or the subset of laminar features (Figure 4 and Table 2). Clustering using laminar features revealed all hippocampal subfields with reasonable accuracy with respect to manually defined ground truth segmentations (except CA4, and CA2 and CA3 were not easily distinguished from one-another). This was expected, since laminar features are one of the key criteria used by histologists to define subfield boundaries (e.g. Duvernoy et al., 2013; Ding & Van Hoesen, 2015). However, when we examined morphological features alone we found unsupervised clusters that closely resembled subfields subiculum, CA1, and a combined CA2 and CA3. Additionally, CA4 was assigned the same clusters as CA1, similar to when clustering was performed on all features combined. This outcome was not expected based on histological data, and provides support for the notion that morphological features capture an independent set of subfield-related structural elements. The observation that morphological feature are sufficient to determine most subfield boundaries holds great promise for future refinement of MRI protocols for subfield delineation given that histological- or laminar-level details are not available in current imaging protocols. Indeed, many of the MR-based subfield segmentation protocols presently available rely on some combination of structural landmarks within or surrounding the hippocampus and indirectly on morphological features (see Yushkevich, Amaral, et al., 2015).

### Anterior-posterior structural variance

Anterior-posterior structural differences in the hippocampus are particularly of interest given the growing body of literature suggesting functional anterior-posterior differences across the hippocampus (e.g. Poppenk et al., 2013; Zeidman & Maguire, 2016; Strange et al., 2014; Plachti et al., 2019). Structural anterior-posterior gradients are difficult to assess using conventional histology given that coronal or sagittal sections are typically out of plane with respect to the different subfields in most of the hippocampal head and tail. This highlights the utility of the unique approach provided by the 3D BigBrain dataset. Figure 6 shows the features gyrification, thickness, and neuronal density across the anterior-posterior axis of the hippocampus. Most notable anterior-posterior differences included differences in most features at the very anterior and posterior extents of the hippocampus. Previous work by (Ding & Van Hoesen, 2015) described the anterior most region – the vertical component of the uncus – as containing modified subfields that were much thinner than their counterparts throughout the rest of the hippocampus, consistent with our observations.

The gyrificaton feature was low in the anterior uncus, high in the remainder of the hippocampal head, and gradually decreased towards the posterior of the hippocampus, most notably in CA1. Qualitatively, we have seen similar trends in gyrification in our previous 7T MRI study (DeKraker et al., 2018) and in other work (Chang et al., 2018), though both of these studies are limited in their ability to detect small gyrifications (i.e. those detected in this study had peak-to-peak distances as low as 2mm). Biophysical models of the development of gyrification suggest a relationship between gyrus size and cortical thickness (Zilles et al., 2013; Striedter et al., 2015), but no systematic anterior-posterior differences in thickness were seen in the present data despite clear decreases in gyrification size towards the posterior. Other structures such as white matter might also constrain gyrification patterns (Striedter et al., 2015), which may additionally have consequences for functional properties of different gyri. For example, (Henderson & Robinson 2014) examined gyrification and structural connectivity in the neocortex and found more unified or modular graph theoretical properties within gyri, as opposed to sulcal regions which were more diffusely connected or hub-like. Similarly, (Plachti et al., 2019; Libby et al., 2012) recently performed parcellation of the hippocampus according to its functional connectivity and observed divisions primarily along the anterior-posterior extent of the hippocampus rather than across subfields (though some proximal-distal clustering was also observed, as in the present study). This functional parcellation likely reflects differences in connectivity across the anterior-posterior extent of hippocampus, and may even relate to modular divisions of function within a given gyrus as proposed in the neocortex by (Henderson & Robinson, 2014).

### Data and resources made available

Alongside this publication we release our detailed manually defined hippocampal subfields, unsupervised clustering results, topological unfolding framework, Equivolume laminar model solutions, and each of the unfolded morphological and laminar features computed here. These resources can be used as templates in other studies, or registration of these features to new data in our unfolded space can be used to guide future subfield segmentation. In addition, we have also made all the code used in this project available via Open Science Framework (https://osf.io/x542s/), and a toolbox for performing hippocampal unfolding, feature extraction, and other useful operations on more general datasets can be found at https://github.com/jordandekraker/Hippunfolding.

### Conclusions

In the current project, we mapped the human hippocampus in detail by combining three methods. First, we used a unique dataset, 3D BigBrain, that contains both histological-level detail and macroscopic 3D spatial context. Second, we imposed a topological unfolding framework to the hippocampus, and third, with this framework we extracted a set of morphological and laminar features, the latter of which have been used prolifically in neocortical characterization and parcellation. Using these methods we highlight three novel empirical observations. First, unsupervised clustering of these features closely resembles classically defined hippocampal subfields. Secondly, despite traditional reliance on laminar features in histology, morphological feature alone are sufficient to closely approximate most hippocampal subfields. Finally, some features such as gyrification in CA1 show, at least qualitatively, subfield-specific anterior-posterior differences that might relate to functional differences observed in the literature. Overall these findings highlight new structural characteristics of the hippocampus, and offer promising avenues for improved delineation and characterization of hippocampal subfields using in-vivo neuroimaging.

## Materials and Methods

### Materials

Histological data used in this study came from the BigBrain dataset bilateral 40um^3^ resolution hippocampal blocks stained for neuronal cell bodies (ftp://bigbrain.loris.ca/BigBrainRelease.2015/3D_ROIs/Hippocampus/) in addition to serial section images before 3D reconstruction at 20um^2^ resolution (ftp://bigbrain.loris.ca/BigBrainRelease.2015/2D_Final_Sections/Coronal/Png/Full_Resolution/) (Amunts et al., 2013). Because of the large file sizes, tracing and application of our unfolding framework were performed on downsampled images (80um isotropic) before upsampling by nearest-neighbour interpolation in the case of labelmaps and linear interpolation in the case of unfolding solutions.

### Manual Tracing

Detailed histological tracing was performed for each hippocampus by a combination of manual tracing and the user-guided computational tools in ITK-SNAP 3.6 (Yushkevich et al., 2006). A general label for hippocampal grey matter (subiculum and CA1-4) was traced first, and this tissue was later manually divided into subfields. Only the laminae which contained stained neuronal cell bodies – stratum pyramidale, oriens, and lucidum – were traced (Figure 1). Stratum radiatum, moleculare, and lacunosum (SRLM) and the alveus were not traced even though they are sometimes considered laminae of the archicortex containing dendrites and axons of pyramidal cells (Duvernoy et al., 2013), because they were not stained by this contrast (though note that some of these strata contain interneurons – see Supplementary Materials section A for discussion).

Subfields segmentation (i.e. the division of archicortical grey matter into distinct subfields) was performed in 3D by rater KF according to the criteria outlined by (Ding & Van Hoesen, 2015). To maximally utilize the histological features available in BigBrain, original 20um images were also consulted every 2mm in order to make use of the highest resolution available. These segmentations included the subiculum and CA1-4, but did not differentiate the regions within the subicular complex due to lack of resolution and since no myelin staining was available. Subfields were traced through the entire length of the hippocampus including the uncus and vertical component of the uncus, in which (Ding & Van Hoesen, 2015) describe modified versions of the same subfields. Because the vertical component of the uncus is very thin, the subfields there were not easily discriminable and so they were partially inferred from neighbouring regions of the hippocampus. All borders were then smoothed by morphological closing and opening to remove small border discrepancies between neighbouring slices.

Structures surrounding the hippocampus were traced only in the regions that border the hippocampus. These labels included medial-temporal lobe neocortex (MTLc) (entorhinal and parahippocampal regions), hippocampal-amygdalar transition area (HATA), and indusium griseum (ind. gris.). HATA borders were clearly discriminable from archicortex by a marked change in density and physical separation from archicortical neurons. Ind. gris. and MTLc borders were less clear, and so they were demarcated using the heuristics used in previous work in MRI. See Supplementary Materials section A for further discussion of these structures.

### Topological unfolding framework

In previous work (DeKraker et al., 2018), we imposed a topological unfolding framework on the hippocampus by solving Laplace’s equation over the domain of the hippocampus under multiple sets of boundary conditions: anterior-posterior, proximal-distal, and laminar. The DG was not unfolded. Although it was easily distinguishable from other subfields by its very high cell density it is topologically disconnected from the rest of the archicortex, and therefore would be out-of-plane (i.e. perpendicular) to our unfolded space (see Figure 1 for visualization). The anterior-posterior and proximal-distal solutions can then be used to index regions of the hippocampus in 2D according to its topology, irrespective of inter-individual differences in gyrifications, rotation, curvature, size, orientation, or position of the hippocampus. This provides implicit registration between hippocampi despite inherently different morphologies. We applied the same approach to the current hippocampal traces (see Figure 2 for illustration). However, note that several minor improvements were made to this code which are detailed in Supplementary Materials section B.

Waehnert et al., noted that neocortical laminae are displaced due to curvature in gyri and sulci, and they propose an ‘Equivolume’ model that captures this feature better than a Laplacian (or equipotential) solution (Waehnert et al., 2014). Their model is motivated by the observation that a given lamina, for example near the pial surface, will stretch at the apex of a gyrus and compress at the depth of a sulcus, causing it to become thinner and thicker in these respective regions and vice versa for laminae at the white matter surface. Thus, we also included an alternative laminar indexing system using the Equivolume model solution obtained from Nighres (Landman et al., 2013). Again, this was performed on the downsampled (80um) traces before upsampling as above. The resulting model had fewer gyrification-related artifacts in laminar profiles so was used for all subsequent laminar analyses. However, some other artifacts were observed under this model solution, likely as a result of the rough texture of the subiculum surface (see Supplementary Materials section C for details).

### Morphological feature extraction

Each morphological feature is illustrated in the top left panel of Figure 3. Thickness estimates were obtained across the unfolded space of the hippocampus as in previous work, that is, by generating and measuring streamlines in 3D across the laminar Laplacian solution obtained from our topological unfolding framework. Curvature estimates were obtained by generating a mid-surface along the hippocampus with the vertices being interpolated xyz coordinates from each unfolded point at a laminar distance of 0.5, which is the midpoint between the inner and outer surface. Smoothing of face normals was applied, and mean curvature was then estimated at each vertex (see Supplementary Materials section B for details). The inner (i.e. adjacent to the SRLM; continuous with the neocortical pial surface) and outer (i.e. adjacent to the alveus, continuous with the neocortical white matter surface) surfaces of the hippocampus were rougher than their mid-surface counterpart due to other features such as subicular ‘islands’ of cell bodies shown in Figure 1. We thus additionally computed curvatures of these surfaces after smoothing as above. Gyrification is typically defined as a ratio outer surface area, for example that of a brain mask over gyrified surface area, in this example including sulcal area (Larsen et al., 2006). Since the hippocampus is an open-ended cortical surface it does not map easily to an outer surface area or to a sphere as in the neocortex, and so our unfolding framework instead maps it to a rectangle. We thus defined gyrification as a ratio of native space surface area over unfolded surface area at each unfolded point.

### Laminar feature extraction

We extracted laminar profiles along the Equivolume laminar solution described above, and then summarized these profiles using the same 10 features consistently used by (Amunts et al., 1999). Briefly, this involved sampling staining intensities (y) along a laminar profile through the cortex, and calculating the mean (Mean(y)). This intensity profile was then treated as a distribution (x), and the mean (Mean(x)) and first 3 moments (SD(x), Skew(x), and Kurt(x)) were calculated. The absolute value of the derivative (Abs.Deriv) of the profile was then calculated (y→y.d), and the same measures (e.g. Mean(y.d), Mean(x.d), etc.) were calculated. These methods are illustrated with corresponding terminology at the top of Figure 3.

There were several methods developed for 3D MRI which we were able to incorporate into this analysis and other differences from the analyses performed by (Amunts et al., 1999). Firstly, we sampled laminar profiles under the 3D Equivolume model that minimizes distortions in laminae due to curvature (as discussed above). Secondly, our laminar sampling was not as dense because of the reduced resolution available in the current data and the fact that the laminae of the archicortex are generally thinner than neocortex. Lastly, we included only laminae containing neuronal cell bodies (as discussed above). Further details on these differences between our methods and those of (Amunts et al., 1999) can be found in Supplementary Materials section B.

### Unsupervised clustering

In order to cluster visually-homologous regions of the feature maps into segments, we applied a scale-space representation employing image pyramids. That is, for each of the features extracted, we smoothed the data in unfolded space with a Gaussian kernel and a Laplacian of Gaussian kernel of sizes sigma=[2,4,8,16,32,64] in order to capture features at various spatial scales. The anterior 10% and posterior 10% of each feature was discarded due to high noise.

All morphological and laminar features from the left and right hippocampi were then reshaped into single vectors, z-scored, and then principal components analysis (PCA) was performed. K-means clustering was then computed on the first 7 components, which explained >1% variance each, with a fixed number of output clusters of k=5 (since manual segmentations contained 5 subfield labels). PCA followed by K-means clustering was ideal for this type of analysis for several reasons: 1) co-linearity among features can be clearly assessed using PCA prior to k-means clustering, 2) clusters were expected to be of comparable sizes, which k-means is biased towards, and 3) the number of clusters is knows a-priori. Clusters were then assigned subfield labels based on highest overlap. Dice overlap scores were calculated (Dice, 1945; Sørensen, 1948) in unfolded space for each subfield (i.e. disregarding thickness), excluding the 10% anterior and posterior edges that were removed due to high noise. We also explored clustering under k=[2,4,8,16,32] in order to determine the consistency of subfield or sensitivity to further subdivisions in the data, which is shown in Supplementary Materials section D. PCA variance explained per component, component loadings and visualization of the first 7 components can be viewed in Figure 5, along with the correlation between each feature.

In order to determine whether subfield clustering could be derived using only laminar features or only morphological features alone, we repeated the above process for the subsets of morphological and laminar features separately. We used the same >1% variance explained threshold to remove PCA ‘noise’ components, which resulted in 5 components in the laminar feature clustering and 3 components in the morphological feature clustering. Morphological features of inner and outer surface textures were excluded since they capture subicular ‘islands’ of cell bodies in stratum lacunosum, which could be considered a laminar feature. This exclusion was also based on the limited value of these two features for any MRI assessment.

### Anterior-posterior variance

One hypothesis that we had based on prior literature was that there may be anterior-posterior differences in some aspects of hippocampal structure. We thus plotted select features of interest across the anterior-posterior axis within each subfield. All features can be seen in a Supplementary Materials section A, where we additionally fit linear trends to the data to determine whether anterior-posterior gradients were present in any subfield. In Figure 6 we focus only on the features mean neuronal density (Mean(y)), thickness, and gyrification which most clearly differed between subfields and are of immediate interest in MRI.

## Acknowledgements

This work is supported by a Canadian Institutes for Health Research Project Grant (CIHR Grant # 366062). J.D. is funded through a Natural Sciences and Engineering Research Council doctoral Canadian Graduate Scholarship (NSERC CGS-D). J.C.L is funded through the Western University Clinical Investigator Program accredited by the Royal College of Physicians and Surgeons of Canada and a Canadian Institutes of Health Research (CIHR) Frederick Banting and Charles Best Canada Graduate Scholarship Doctoral Award. K.M.F is funded through an Ontario Graduate Scholarship (OGS).

We would also like to thank Dr. Alan Evans, Dr. Katrin Amunts, and all contributors to project 3D BigBrain for developing and releasing this invaluable resource.

