## Supplementary Materials for "Hippocampal subfields revealed through unfolding and unsupervised clustering of laminar and morphological features in 3D BigBrain"

### Section A: additional anatomical details of the hippocampus and surrounding structures

The dentate gyrus, modelled in Figure 1A, was excluded from our hippocampal unfolding framework because it was used as a boundary condition for proximal-distal coordinates. However, it was easily differentiated from other subfields by its very high neuronal density. Figure 1B shows residual staining at tissue boundaries, as well as some staining within the SRLM laminae which may be due to the presence of inhibitory interneurons.

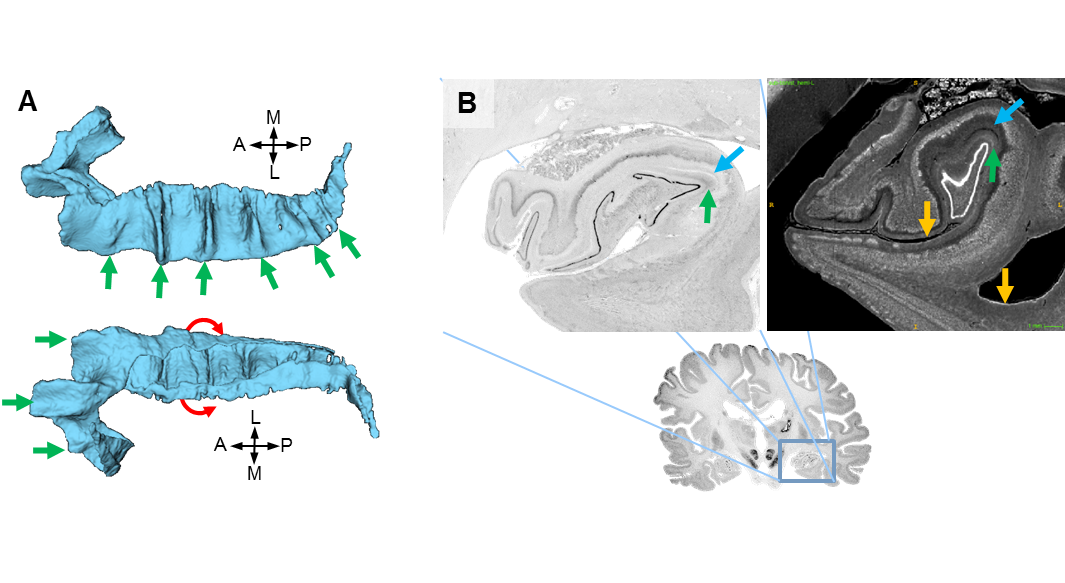

Figure 1. Anatomical details noted in BigBrain 3D histology outside of ‘archicortex’ label. A) 3D models of the dentate gyrus from the superior (top) and inferior (bottom), with the red line demonstrating the ‘U’ shape and green arrows indicating ‘dentations’ that are prominent in human DG and follow the gyrification of archicortex. B) Increased staining can be seen on CSF boundaries (e.g. orange arrows) but also may include interneurons found in the stratum moleculare of the CA fields (blue arrows) and stratum moleculare of the dentate gyrus (green arrows), which are intermittently fused or separated by the vestigial hippocampal sulcus. Images shown are from the original 20um resolution slices (left) and a coronal slice from the 40um isoluminant hippocampal block (right).

We detected gyral peaks in the left and right hippocampi, and calculated the distance between them. Briefly, peaks were detected by taking an anterior-posterior profile midway through manually defined subfield CA1 (where most gyri were centered) and detecting local maxima in curvature (see sections 2.3 and 2.4). Distances between gyral peaks are shown in Figure 2, and were generally low in the uncus, high in the remaining hippocampal head and anterior body, and decreasing in size through the body and tail of the hippocampus. Smaller gyri had as little as 0.4mm of tissue (alveus on the outer surface or SRLM on the inner surface) separating them.

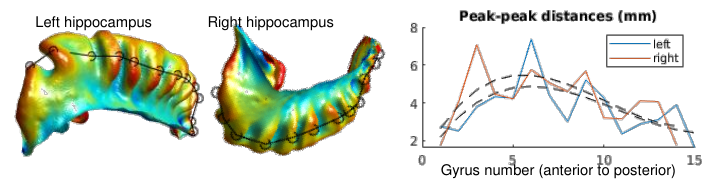

Figure 2. Peak-peak distances between hippocampal gyri.

Figure 3 shows the distributions of all these data from both the left and right hippocampi, colour-coded by subfield. Linear trends where R-squared values were greater than 0.1 are overlaid. Laminar feature Mean(y), or the mean amount of staining, increased towards the posterior of the hippocampus in regions CA1 and CA4. Mean(x.d), or the depth at which the greatest change in staining intensity was seen, increased in CA2 and CA3 towards the posterior as well. Skew(x), or the skew on the depth of neurons, decreased towards posterior subiculum. The morphological feature gyrification also showed a decrease towards the posterior in most prominently in CA1 which was also noted in Figure 1, but also in CA3.

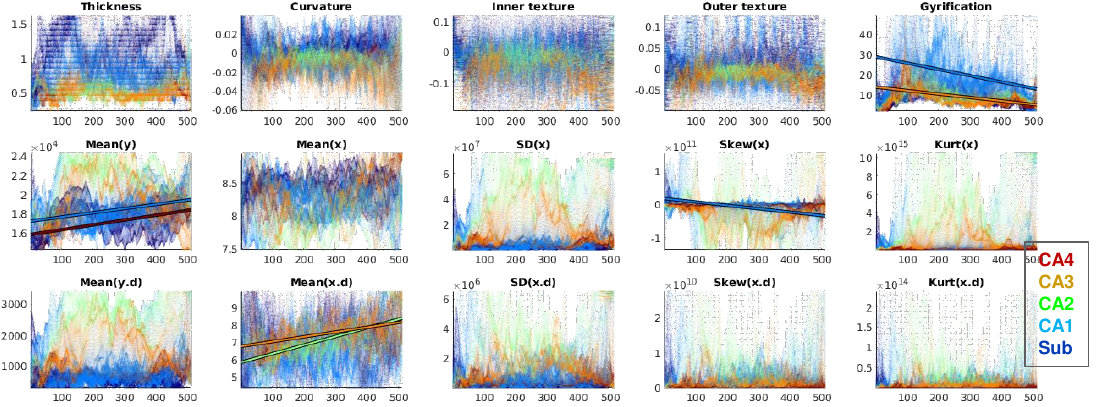

<2 column width> Figure 3. Examination of anterior-posterior differences in hippocampal structure. The x-axes represents the anterior-posterior axis of the hippocampus and the y-axes are the corresponding feature values, colour-coded by manual subfields. Linear correlations with R-squared values greater than 0.1 are overlaid.

### Section B: additional methodological details

Our Laplacian unfolding framework was modified in several ways since its use in our previous publication (DeKraker et al. 2018):

1) DG was excluded from the domain of hippocampal archicortex (though CA4 was instead differentiated from CA3).
2) Unfolded representations were modified to have a 2:1 rather than 1:1 anterior-posterior:proximal-distal aspect ratio which we found to more closely match the real world dimensions of hippocampal tissue. BigBrain flatmaps were generated with 512x256 coordinate points in order to capture the higher level of detail available in that dataset (previous work was 100x100 points).
3) General code optimizations to make it faster given large datasets and more robust to errors in manual segmentation. Interpolated mid-surfaces are now also automatically generated. These changes do not affect the unfolding outputs.

Updated code and the history of changes made can be seen at https://github.com/jordandekraker/HippUnfolding.

The Equivolume model solution was obtained by a python wrapper of ‘CBStools’ called ‘Nighres’ (https://github.com/nighres/nighres).

Laminar sampling differences between Amunts et al. and the current study differed in the following way: horizontal (i.e. perpendicular to each profile) smoothing was performed with a Gaussian kernel of sigma=3 in unfolded space, rather than with a parametrically optimized sigma in native space (sigma could not be optimized to detect 3-5 laminar profile peaks in the neocortex since we expected only 1-3 laminar profile peaks in the archicortex and we wanted to avoid overfitting).

Mean curvature was estimated using the function ‘patchcurvature()’ from the MathWorks File Exchange (https://www.mathworks.com/matlabcentral/fileexchange/32573-patch-curvature).

### Section C: laminar modelling using Laplacian vs. Equivolume model

The Equivolume model aims to account for displacements of laminae due to curvature. The Laplacian model remains smooth even over complex shapes, but does not account for displacement of laminae due to curvature, and so there is an offset between the depths at which peak staining along a laminar profile is found. The Equivolume model more closely aligns peaks between gyral and sulcal areas, but is highly susceptible to distortion from the rough surfaces of detailed manual segmentations (e.g. due to subicular ‘islands’), which can be seen in the sagittal slice of the model solution.

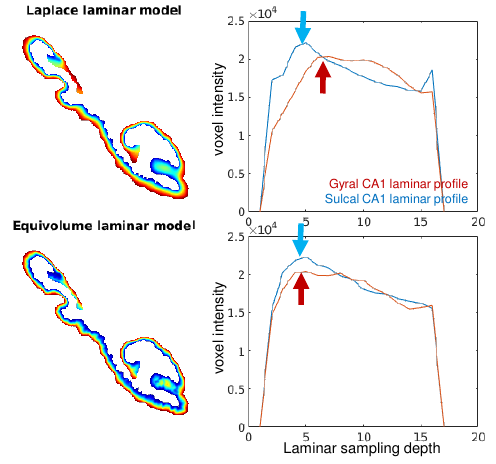

Figure 4. Comparison of Equivolume and Laplacian models of archicortical lamination. The left images show a sagittal slice of each model solution in the right hemisphere. Right images show two example laminar profiles, one at the peak of a gyrus (A-P,P-D coordinates 236,104) and one at the depth of sulcus (A-P,P-D coordinates 218,104) in the CA1 of the right hippocampus.

### Section D: additional k-means clustering results

By setting k=5 in k-means clustering we imposed some prior information onto the segmentation of the hippocampus. Thus, we also explored different possible parcellation schemes by setting k=[2,4,8,16,32]. We ordered the resulting clusters according to their median proximal-distal distance in order to better align with our manual segmentation label scheme. Within each clustering result, borders can be seen near the manually defined subfield borders. With greater numbers of clusters, the subfields divide into additional regions but most borders separate proximal-distal regions rather than anterior-posterior regions. This could be due to the presence of transition zones between subfields, or perhaps in the case of subiculum additional proximal-distal segments become clustered due to subicular subregions (e.g. prosubiculum, presubiculum, subiculum proper, or parasubiculum). Overall these data provide strong motivation for the segmentation of the hippocampus along its proximal-distal rather than its anterior-posterior axis, and the most seen borders resemble the classic hippocampal subfield definitions.

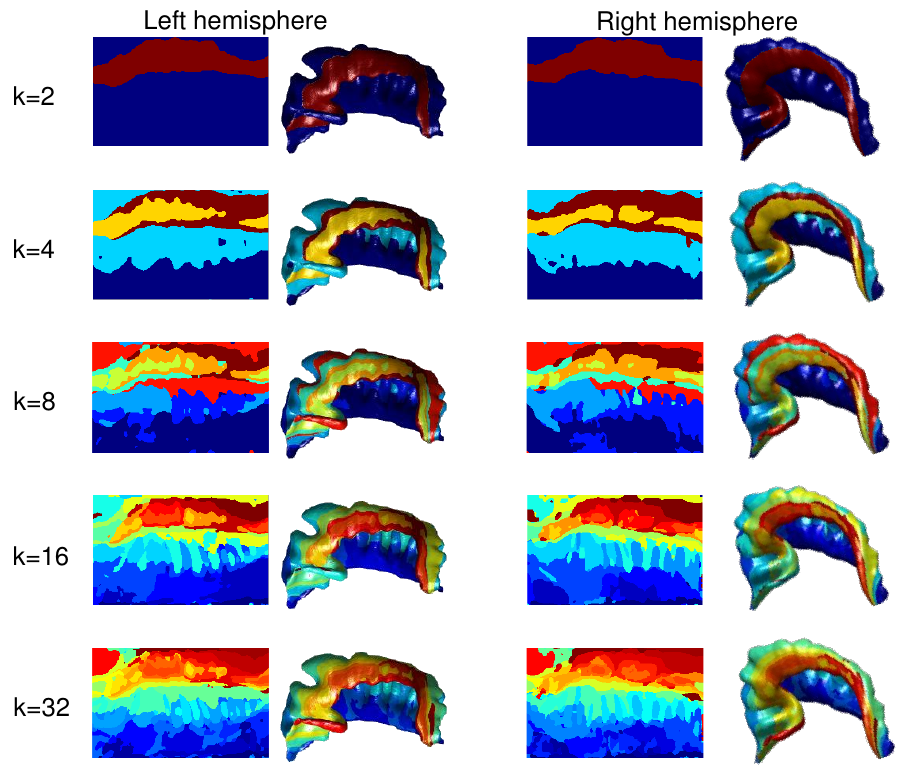

Figure 5. Exploration of different clustering schemes by varying k in k-means clustering. Resulting clusters are labeled and coloured according to their median proximal-distal distance.
